# Using software Vortex as a tool towards conservation actions for sharks

**DOI:** 10.1101/831677

**Authors:** Laís Ramos Barcellos, Fabricio Escarlate-Tavares

**Affiliations:** Centro Universitário de Brasília, SEPN 707/907, Brasília, DF, Brasil

**Keywords:** Elasmobranchs, Population Viability Analysis, Vortex, Demography

## Abstract

Elasmobranchs, especially sharks, are part of the most threatened group of animals in the world. Among the factors that negatively affect these populations, the environmental degradation and disordered fishing can be highlighted. Although these threats are well known, the responses of most species to these factors are not fully understood, which makes it difficult to predict potential population responses according to conservation actions. The present study aims to develop a theoretical model based on a real population of white sharks (*Carcharodon carcharias*) that inhabit Gulf of Mexico, using their biological patterns to identify trends for the populations using Vortex (version 10.0.0.3) as a tool. The construction of the model was based on bibliographic data, considering parameters of life history, demographic rates and main known threats. Data were entered into Vortex software, and two population sizes were tested in the population viability analysis, indicating in both cases that the species tends to lose its genetic diversity over 500 years with a 12% and 51% of probability of extinction, respectively, considering exclusively its biological potential. This results can reflect on an mislead assumption of the real subpopulation’s size, which shows that the species is inserted into the vortex of extinction even with the existence of protective measures regarding white sharks on the location of the study, and it must incorporate neonates and juveniles individuals on the analysis to generate a consistent result. Despite this, the study suggests that the utilization of Vortex is capable of generating responses necessary for the conservation of sharks by analyzing stochastic events and life history in a simple way.

## 1. INTRODUCTION

Sharks are apex predators in the food chain in different marine ecosystems; however, they are susceptible to decline in all the oceanic basins, turning the state of conservation of many populations a considerable concern (Myers, 2007). According to Evans (2001), sharks have a vital role in the maintenance of the ecological balance in the oceans, and, in this context, the establishment of adequate measures to the management of marine elasmobranchs are fundamental for the health of many marine ecosystems.

The main treat to elasmobranchs is uncontrolled fishing (Bornatowski, 2012). The capture of these animals has been increasing globally, and it is estimated that about one million tons per year were captured in the last decades (Bonfil, 1994). Different methods of modern fisheries are exploring a wide variety of sharks for human consumption and also to commercialize their fins (practice internationally known as finning). There is still the accidental catch when fishing for other species, threat known as by-catch (Musick, 2005). However, the management of elasmobranch stocks is hindered mainly by the lack of basic information on the dynamics of populations worldwide (SBEEL, 2009).

Some of the group’s biological characteristics result in intrinsic growth rates and very low resilience capacity, as well as slow growth, late sexual maturity and low fertility, and these conditions cause populations to be more susceptible to overfishing due to the low replacement of new individuals, which leads to the decline in populations (Hoenig et al., 1990; Sminkey & Musick, 1995, 1996; Camhi et al., 1998; Musick, 1999; Smith et al., 1998). There is a concern about sustainability by the International Union for Conservation of Nature (IUCN), because sharks are not specifically managed or conserved by any regional or multilateral treaty that aims the management of marine resources (IUCN, 2002).

The biology of most species of sharks is little known and there are poor data about demography and life history (Pierce et al, 2009; Frisk et al, 2010). Information about their population demography offers ways to estimate intrinsic population growth rate “r” in response to impacts as fishing, and also allows to predict the future tendencies for their population size according to the life history of the analyzed species (Gilpin et al., 1986; Caillet et al., 2005; Dulvy & Forrest, 2010). With these data combined with studies on population structure, age and growth and distribution, it can be possible to provide a proper management to each species (Palsboll et al., 2007).

The Population Viability Analysis (PVA) models the effect of deterministic processes, inbreeding, allelic drift, environmental and demographic stochasticity in a population (Lacy, 2005). This analysis predicts the tendencies in population size to increase or decrease in the future according to the life history of the species analyzed and the current situation of the population, taking into consideration the deterministic or stochastic factors that may cause fluctuations in the population size (Gilpin et al.,1986). However, its widespread use is hampered by the scarcity of information on vital rates and lack of ecological information of the species, which are required as input parameters for the models (Pardini, 2001; Beerkircher, 2003).

The PVA assesses the risk of a population of wildlife animals to diminish or extinguish due to current or future conditions. Demographic information of the population serve as data to be added to a “Vortex” model with a structure that defines the basic biological characteristics of the species and habitat use patterns, so the model can project the demographic behavior of the simulated population for a specific period of the future, considering the specific conditions assumed. Within these aspects, the preconditions for growth or decline of the population can be determined, as well as the best options for management of the species in order to minimize the risk of extinction (Lacy, 1993).

This study aimed to compile biological and population data of white sharks *Carcharodon carcharias* (Linnaeus, 1758) through literature review, since this species has lots of biology and life history data available of the population that inhabits the northeastern Pacific Ocean based on a theoretical but biologically accurate population tested in different scenarios to analyze if using Vortex as a tool is applicable to identify future shark’s population trends.

## 2. METODOLOGY

### 2.1 BIBLIOGRAPHIC RESEARCH

The construction of the theoretical model was based on bibliographic search considering life history parameters, demographic and biological rates, and threats known from different populations. The individuals that inhabit the northeast Pacific Ocean were chosen for the analysis due to almost all individuals were properly registered, based on the article by Chapple et. al. (2011) and Burgess et al. (2014), therefore the population size was known.

A variety of sources were used to improve quantitative and qualitative information and to develop the estimates used in this analysis. Scholar Google, Research Gate and Scielo were used as primarily data survey for the PVA. The purpose of the data survey was to gather as much information as possible about the biology of the species, and it were not established a data limit, number of articles or quotes.

Some key-words were detected as relevant for the bibliographic search to evaluate the population viability: great white sharks, ICCAT, FAO, IUCN, CITES, population dynamics, mortality, sexual maturity, gestation, *Carcharodon carcharias*, biology, phylogenetic, extinction, capture, reproduction, inbreeding, shark population model, shark demographic modeling, Vortex software.

### 2.2 MODELLING

The data obtained from the literature were condensed in Table 1 and then inserted into the software Vortex (version 10.0.0.3), which uses the Monte Carlo simulation that can determine the independent variables of possible values and behaviors related to them, and also describes the distribution and characteristics of possible values of a dependent variable. This simulation is used to incorporate uncertainty in the parameters and generate demographic population growth rates, generation time and elasticity for a large set of matrices which includes several parameters (Miller & Lacy, 2005).

**Table 1.**
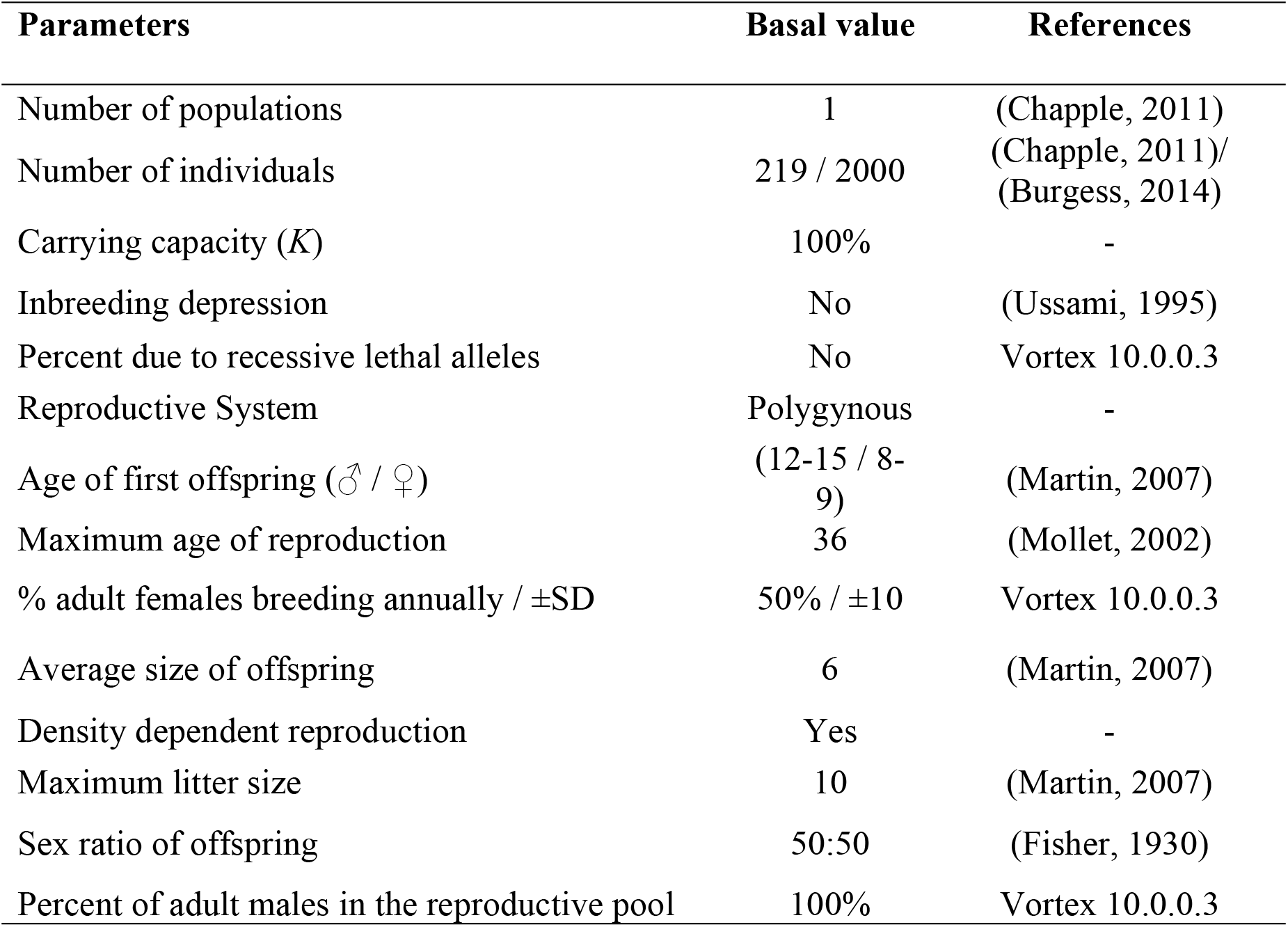
Synthesis of data used in the PVA.

| Parameters | Basal value | References |
| --- | --- | --- |
| Number of populations | 1 | (Chapple, 2011) |
| Number of individuals | 219 / 2000 | (Chapple, 2011)/<br>(Burgess, 2014) |
| Carrying capacity (*K*) | 100% | - |
| Inbreeding depression | No | (Ussami, 1995) |
| Percent due to recessive lethal alleles | No | Vortex 10.0.0.3 |
| Reproductive System | Polygynous | - |
| Age of first offspring (♂ / ♀) | (12-15 / 8-9) | (Martin, 2007) |
| Maximum age of reproduction | 36 | (Mollet, 2002) |
| % adult females breeding annually / ±SD | 50% / ±10 | Vortex 10.0.0.3 |
| Average size of offspring | 6 | (Martin, 2007) |
| Density dependent reproduction | Yes | - |
| Maximum litter size | 10 | (Martin, 2007) |
| Sex ratio of offspring | 50:50 | (Fisher, 1930) |
| Percent of adult males in the reproductive pool | 100% | Vortex 10.0.0.3 |

The basal model was defined to investigate the PVA of white sharks from the northeast of the Pacific Ocean based on this theoretical but biologically accurate population. The name given to the initial tested population was “Northeast of the Pacific Ocean”. As the longevity of the specie is 36 years old (Mollet, 2002), there were analyzed interactions in the period of 500 years to better observe the tendency of population to grow or decrease during their generations.

According to Ussami (1995), it was considered that there is no inbreeding depression. Little is known about the mating system of white sharks, however, giving into consideration the life history and that these animals are solitary, we assumed that they are polygamies. According to Mullet (2002), they meet in the period of two or three years to breed, with gestational duration of 12 months. In the present study, it was considered that the reproduction depends on the density based on biological aspects, because these animals are solitary and depend on the meeting between individuals to reproduce.

According to Martin (2007), males reproduce for the first time between ages 12 and 15, and females between 8 and 9. In Simulation 1, it was considered that females have their first offspring at 8 years old, and males copulate for the first time at 13. In Simulation 2, it was considered that females reproduce with 9 years old and males at 15.

According to Preston (1995), the infant mortality rate can reach over 80%. For adults, Baum et. al (2002) estimated that for every 1000 hooks, two white sharks are caught in the Gulf of Mexico. Using the population parameters of Mollet (2002), the mortality rate (*M*) is 0.153, or 86% chance of survival.

In the tested basal models, mortality rates were estimated automatically by the software, having Table 2 as a synthesis of the rates for males and females.

**Table 2.** Annual mortality rates of white sharks according to Vortex software.

| **Mortality rate (*M*) according to age** | ♀ | ♂ |
| --- | --- | --- |
| 0 to 1 year (SD ± 10) | 50% | 50% |
| 1 to 36 years (SD ± 3) | 10% | 10% |

Therewith, two simulations were analyzed, changing population sizes and age at first reproduction among them. According to Chapple et. al. (2011), the initial population that inhabits the northeastern Pacific Ocean consists of 219 individuals, with 106 individuals with known sex. However, Burgess et. al (2014) in his review to Chapple’s article concluded that at least 2000 individuals were present at the Central California region.

The support capacity was considered as 100, taking advantage of the result that the software itself provided. None environmental variation was added in the support capacity and none catastrophe as well. No supplementation deriving from other unrelated populations was incorporated into the demographic model.

## 3. RESULTS

The software estimated that according to the previous data patterns and concepts of life histories of the animals, 50% of the population reproduces at low density, and 25% reproduces in the supporting capacity (K=100). It was considered that all adult males were part of the reproductive pool, 65.7% of this generating brood, having the value of 1.6 as an average of successful mating.

Only one population was considered in the basal model, since it was analyzed only the subpopulation that inhabits the northeastern Pacific Ocean, with immigration or emigration.

### 3.1 Sexual Ratio

According to Chappel et. al. (2011), the sexual ratio taking into consideration the minimum value of abundance (69 males; 19 females; 42 unknown), is probably biased for males because it is easier to confirm the presence of claspers than absence of it, leading to larger numbers of unknown sex. Chapple also considers in his study that there are only sub adult and adult individuals in the studied population.

In Simulation 1, it was considered the age distribution according to the article by Chapple et. al. (2011), where it is estimated that the subpopulation of northeastern Pacific Ocean comprises 219 individuals. The total population used in the models was 106 individuals, as there are certainly 73 males and 33 females composing this population, ignoring the individuals where sex is unknown.

In Simulation 2, it was considered 2000 individuals composed the subpopulation. Thus, it was possible to synthesize sizes regarding age and the number of individuals by gender (Table 3), using the estimation of the age composition of white sharks according to the mark-recapture study by Chapple et al. (2011) combined with the theoretical age distribution by Cailliet et al. (1985). As there is a lack of record for several age groups, none individual composed neonate and juvenile slots.

**Table 3.** Estimation of the age composition of white sharks according to the mark and recapture study by Chapple et. al. (2011) combined with the theoretical age distribution by Cailliet et. al. (1985).

| **Length (cm)** | **Age related to growth** | **106 individuals**<br>**(CHAPPLE, 2011)**<br>**(♂ / ♀)** | **2000 individuals**<br>**(BURGESS, 2014)**<br>**(♂ / ♀)** |
| --- | --- | --- | --- |
| 245 | 4 | 0 / 0 | 0 / 0 |
| 275 | 5 | 0 / 0 | 0 / 0 |
| 305 | 6 | 0 / 2 | 0 / 38 |
| 335 | 7 | 1 / 0 | 19 / 0 |
| 365 | 8 | 7 / 4 | 132 / 75 |
| 395 | 10 | 9 / 2 | 170 / 38 |
| 425 | 11 | 16 / 4 | 302 / 76 |
| 455 | 13 | 22 / 3 | 415 / 57 |
| 485 | 15 | 14 / 11 | 264 / 207 |
| 515 | 17 | 4 / 7 | 75 / 132 |
| 545 | 20 | 0 / 0 | 0 / 0 |

### 3.2 Modelling

Table 4 shows the results of the two simulations inputs on the software Vortex, which each differed according to the population size, first age reproduction and number of interactions to be analyzed. With (r) being the stochastic rate of population growth; SD (r) being the standard deviation of stochastic population growth rate; P.E. being the Probability of Extinction; Div. Gene being the Genetic Diversity and N being the population’s size final mean.

**Table 4.** Result’s summary of the two simulations modeling the subpopulation of northeastern Pacific Ocean without dispersion.

| Population of northeastern Pacific Ocean |  |  |  |  |  |
| --- | --- | --- | --- | --- | --- |
|  | r | SD (r) | P.E. | Div. Gene | N |
| Simulation 1 | 0,006 | 0,099 | 0,12 | 51% | 65 |
| Simulation 2 | -0,006 | 0,104 | 0,57 | 43% | 51 |

### 3.3 Simulation 1

The data of population size to be analyzed was inserted in the software, comprising 106 individuals having 500 independent interactions over 500 years. It was considered that females have their first offspring at age 8 old and males copulate for the first time at age 13 (Figure 1).

**Figure 1.**
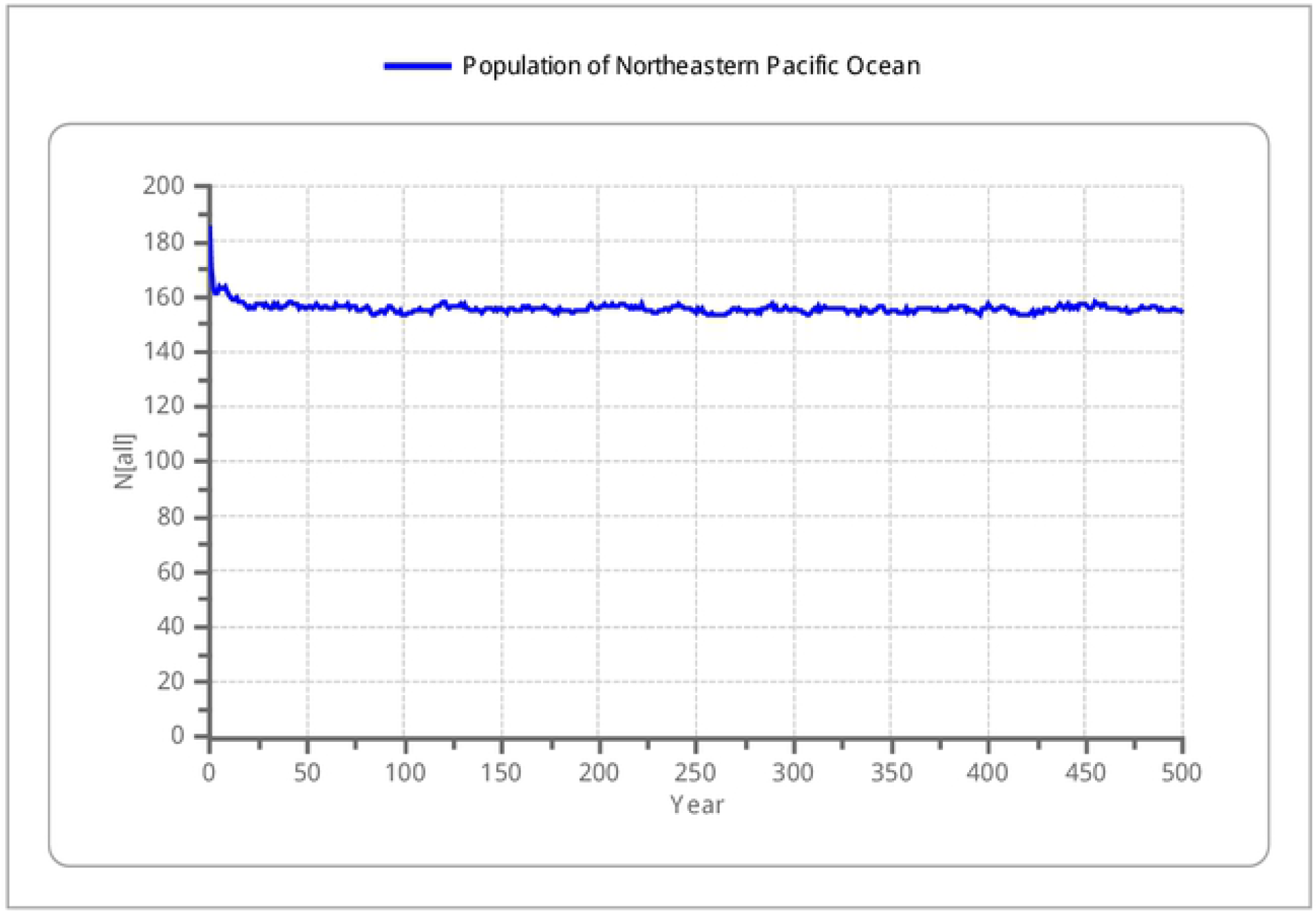
Representation of the results of Simulation 1.

It can be observed that the trend of this population is to decrease over 500 years, with a 12% extinction probability, leaving an average of 65 individuals and being stabilized after 25 years, which is considered to be inserted into the vortex of extinction, since the loss of genetic diversity is nearly 50%, thereby undergoing to the genetic bottleneck.

### 3.4 Simulation 2

For this simulation, it was considered a total of 2000 individuals, 1000 interactions in 500 years, with females producing their first offspring at age 9 and males reproducing for the first time at age 15 (Figure 2).

**Figure 2.**
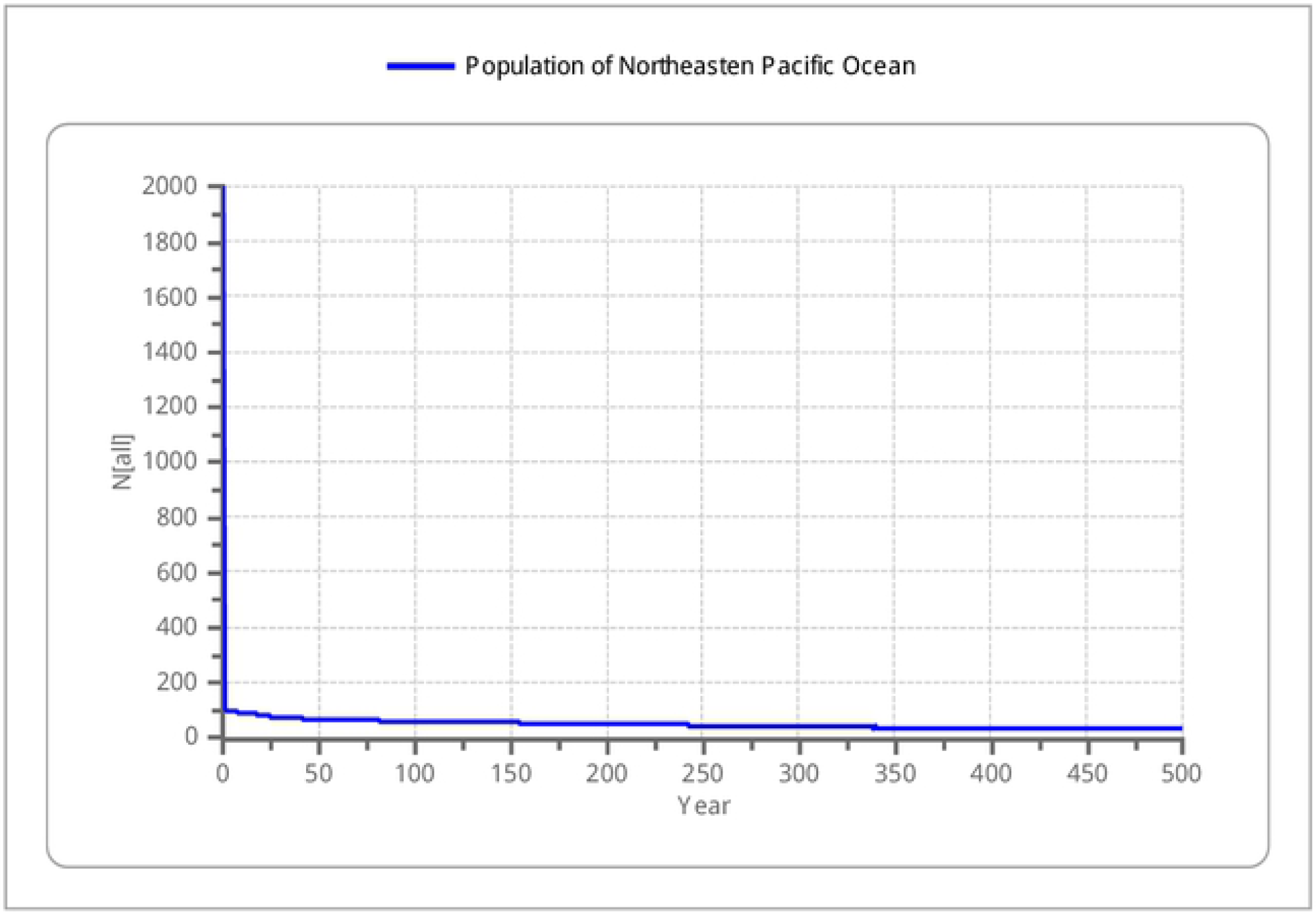
Representation of Simulation 2.

The population growth rate is negative, there is an abrupt drop in the first 5 years and it is assumed that this is related to the age distribution from Table 4, because an extrapolation of the proportion based on the mark-recapture study by Chapple et al. (2011) was used for the population estimation by Burgess et al. (2014), and this lead the probability of extinction to reach 57%, leaving an average of 51 individuals at the end of 500 years, which indicates that the population is inserted into the extinction vortex with high loss of genetic diversity through a drastic reduction in the population size.

## 4. DISCUSSION

White sharks are highly sensitive to over-exploitation and are currently in the IUCN Red List as Vulnerable. These animals are globally distributed in at least three genetically distinct populations, which are located in South Africa (<u>Pardini</u> *et al*., <u>2001</u>; <u>Bonfil</u> *et al*., <u>2005</u>), Australia/New Zealand (<u>Pardini</u> *et al*. <u>2001</u>; <u>Bruce</u> *et al*., <u>2006</u>) and the northeastern Pacific Ocean (<u>Boustany</u> *et al*., <u>2002</u>; <u>Weng</u> *et al*., <u>2007</u>; <u>Domeier & Nasby-Lucas, 2008</u>), the last one migrating from Guadalupe Island in Mexico to the coast of California in the United States (Chapple et al., 2011).

It is the first time that the software Vortex is used for PVA of elasmobranchs, and since the software is mostly used for birds and mammals, the hypothesis was that it might work for sharks is because they have biological peculiarities that bring them much closer to reptiles and birds in terms of vulnerability than to bony fish themselves (Hoenig & Gruber, 1990).

Both models are not considered a real tendency on the sub-population of Northeast Pacific because of the extrapolating number of adults and sub-adults composing the population and zero presence of neonates or juveniles, which is an answer to why the value of intrinsic growth rate in Simulation 2 is demonstrating a decline. It is known from the study by Cortés (2007) that juveniles presence in populations have the most influence on population growth rate, and assuming that there are no presence of this age classes is probably an error in the tag-recapture study by Chapple et al. (2011).

Also, the two models from the present study have a total population size of <100, which definitely leads to a loss of genetic variability and high extinction rates according to Blower et al. (2012), that suggests by genetic studies that a population require a minimum of 500-1000 breeding individuals to not undergo to the genetic bottleneck. The demographic randomness varies with time, and when the genetic characteristics of the population are considered, it can generate low population rates and loss of genetic variability, causing reduction of allelic variation (Guedes, 2004).

Vortex was developed with the purpose of modeling based on one or more known populations of a certain species and their future tendencies. By considering their biological parameters, the data is submitted to different treatments and allow the knowledge of the process that contribute to the vulnerability of the population and evaluate alternatives to improve the conservation of the species (Lacy, 1993). However, there are failures when using Vortex due to not take into consideration the process of natural variances, as well as uncertainty in the vital estimation of species. Despite this, the software is useful because it provides a simple population growth model and captures the essential information of changes in population size, enabling to make predictions about the future of this population, which assists significantly in decision-making for strategies focused on the conservation of species.

This software, as well as other methods of modelling, is not intended to give definitive answers, but to indicate tendencies. It is important to consider the effects that the uncertainty and the variation in vital rates may have on population parameters, specially referring to marine species, where the vital rates estimative are difficult to obtain and often result in great uncertainty (Caswell, 1998).

The software has three options to analyze the tendencies of extinction in years, being 100, 500 or 1000 years. Considering that the specie’s longevity is 36 years old, if 100 years were chosen, only three generations would be observed. Besides the results from this study, there are signs that this sub-population has increased the number of juveniles, showing that at least the efforts on protection are working (Lowe et al., 2012).

## 5. CONCLUSION

The utilization of PVA is growing increasingly in the conservation context. This tool is capable of generating responses necessary for the conservation of the species by analyzing stochastic events and life history in a simple way. This study aimed to introduce a new method on the conservation of sharks, and is not a real representation on the future of white sharks.

A more detailed study with representative data on real population size is needed. Pursuant to the results shown in the present study, the tendency of diversity genetic loss of the population analyzed reaches 50%, a result reflected on an mislead assumption of the real subpopulation’s size, which shows that the species is inserted into the vortex of extinction even with the existence of protective measures regarding this species on the location of the study.

## AKNOWLEDGEMENTS

We thank Centro Universitário de Brasília (UniCEUB) for the support, and Rogério Cunha de Paula for the contributions.

## REFERENCES

Baum JK, Myers RA, Kehler D, Gerber L, Blanchard W, Harley SJ. Preliminary standardized catch rates for pelagic and large coastal sharks from logbook and observer data from the northwest Atlantic. Coll Vol Sci Papers ICCAT. 2002;54(4): 1294–1313.

Beerkircher LR, Shivji MS, Cortés E. A Monte Carlo demographic analysis of the silky shark (Carcharhinus falciformis): implications of gear selectivity. Fish Bull. 2003;101: 168–174.

Blower DC, Pandolfi JM, Bruce BD, Gomez-cabrera MC, Ovenden JR (2012). Population genetics of Australian white sharks reveals fine-scale spatial structure, transoceanic dispersal events and low effective population sizes. Mar Ecol Progr Series. 2012;455: 229–244. doi: 10.3354/meps09659.

Bonfil R. Overview of world elasmobranch fisheries. FAO Fish Tech Paper, Rome. 1994;341: 119.

Bonfil R, Meyer M, Scholl M, Johnson R, O’brien S, Oosthuizen H, Swanson S, Kotze D, Paterson M. Transoceanic migration, spatial dynamics, and population linkages of white sharks. Science. 2005;310: 100–103. doi: 10.1126/science.1114898.

Bornatowski H, Abilhoa V. Tubarões e raias capturados pela pesca artesanal no Paraná: guia de identificação. Hori Cad Téc. 2012;4, 124.

Boustany AM, Davis S, Pyle P, Anderson S, Leboeuf B, Block B. Expanded niche for white sharks. Nature. 2002;415: 35–36.

Burgess GH, Bruce BD, Cailliet GM, Goldman KJ, Grubbs RD, Lowe CG, Macneil MA, Mollet HF, Weng KC, O’sullivan JB. A re-evaluation of the size of the White shark (Carcharodon carcharias) population off California, USA. Plos One. 2014; 9: 6. doi:10.1371/journal.pone.0098078.

Cailliet GM, Natanson LJ, Welden BA, Ebert DA. Preliminary studies on the age and growth of the white shark, Carcharodon carcharias, using vertebral bands. South Cali Acad of Sci. 1985; 9: 49–60.

Cailliet GM, Musick JA, Simpfendorfer CA, Stevens JD (2005). Ecology and life history characteristics of chondrichthyan fish. In: Fowler SL, Cavanagh RD, Camhi M, Burgess GH, Cailliet G, Fordham SV, Simpfendorfer CA, Musick JA, editors. Sharks, Rays and Chimaeras: The Status of the Chondrichthyan Fishes, Status Survey. IUCN: Gland, Switzerland and Cambridge, UK; 2005. pp.12–18.

Camhi M, Fowler SL, Musick JA, Bräutigam A, Fordham SV. Sharks and their Relatives – Ecology and Conservation. IUCN/SSC Shark Specialist Group. IUCN, Gland, Switzerland and Cambridge, UK; 1998.

Caswell H, Brault S, Read AJ, Smith TD. Harbor porpoise and fisheries: an uncertainty analysis of incidental mortality. Ecol Applic. 1998;8(4): 1226–1238.

Chapple TK, Jorgensen SJ, Anderson SD, Kanive PE, Klimley AP, Botsford LW, Block BA. A first estimate of white shark, Carcharodon carcharias, abundance off Central California. Biol Lett. 2011;7(4): 581–583. doi:10.1098/rsbl.2011.0124.

Cortés E. Chondrichthyan demographic modeling: an essay on its use, abuse and future. Mar Fresh Res. 2007;58: 4–6.

Domeier M, Nasby-lucas N. Migration patterns of white sharks Carcharodon carcharias tagged at Guadalupe Island, Mexico, and identification of an eastern Pacific shared offshore foraging area. Mar Ecol Prog Series. 2008;370: 221–237. doi: 10.3354/meps07628.

Dulvy NK, Forrest RE. Life histories, population dynamics, and extinction risks in chondrichthyans. In: Carrier JC, Musick JA, Heithaus, MR, editors. Sharks and their relatives II: biodiversity, adaptive physiology, and conservation. Boca Raton: CRC Press; 2010. pp.635–676.

Evans MD. Shark Conservation: The Need for Increased Efforts to Protect Shark Populations in the Twenty-First Century. Penn St Environ Law Rev. 2001; 10(1): 21.

Fisher RA. The Genetical Theory of Natural Selection. Clarendon: Oxford; 1930.

Gilpin ME, Soulé ME Minimum viable populations: processes of species extinction. In: Soulé ME, editor. Conservation Biology: The Science of Scarcity and Diversity. Sunderland, MA; 1986. pp. 19–34.

Guedes FB. Genética da conservação como uma ferramenta para avaliar os problemas populacionais da fragmentação de habitat. Monography (Graduation), Setor de Ciências Biológicas da Universidade Federal do Paraná. 2004. Available from: https://www.acervodigital.ufpr.br/bitstream/handle/1884/32965/MONOGRAFIA%20FATIMA%20BECKER%20GUEDES.pdf?sequence=1

Hoenig JM, Gruber SH. Elasmobranchs as living resources: advances in the biology, ecology, systematics, and the status of the fisheries. In: Pratt HL, Gruber SH, Taniuchi T, editors. Life-history patterns in the elasmobranchs: implications for fisheries management. NOAA/National Marine Fisheries Service; 1990. pp. 1–16.

IUCN. Species Survival Commission’s Shark Specialist Group and TRAFFIC, The role of CITES in the conservation and management of Sharks. 2002. Available from: http://www.cites.org/common/notif/2002/ESF042A.pdf.

Lacy RC. VORTEX: A computer simulation model for Population Viability Analysis. Wild Res. 1993;20: 45–65.

Lacy RC, Miller PS. VORTEX. A stochastic simulation of the simulation process. Version 9.50 user’s manual. Conservation Breeding Specialist Group (IUCN/SSC). 2005. Available from: http://www.vortex10.org/Downloads/v950Manual.pdf.

Lowe CG, Blasius ME, Jarvis ET, Mason TJ, Goodmanlowe GD, O’sullivan JB. Historic fishery interactions with white sharks in the southern California Bight. In: Domeier M, editor. Global perspectives on the biology and life history of the great white shark. Boca Raton: CRC Press; 2012. pp. 169–185.

Martin JB. The price of fame: Cities regulations and efforts towards international protection of the great white shark. The Geo Wash Intern Rev. 2007;39(1), 199.

Mollet HF, Cailliet GM. Comparative population demography of elasmobranchs using life history tables, Leslie matrices and stage-based matrix models. Mar Fresh Res. 2002;53(2): 503–516. doi: 10.1071/MF01083.

Musick JA. Life in the slow lane: Ecology and conservation of long-lived marine animals. Amer Fish Soc Symp. 1999; 23: 1–10.

Musick JA, Ellis JK. Reproductive Evolution of Chondrichthyes. In: Hamlett WC, editor. Reproductive Biology and Phylogeny of Chondrichthyes: Sharks, Batoids and Chimaeras. Science Publishers, Inc. Plymouth, UK; 2005. pp. 45–79.

Myers RA, Baum JK, Shepherd TD, Powers SP, Peterson CH. Cascading effects of the loss of apex predatory sharks from a coastal ocean. Science. 2007;315: 1846–1850. doi: 10.1126/science.1138657.

Palsboll P, Berube M, Allendorf F. Identification of management units using population genetic data. Tren Ecol Evol. 2007;22: 11–16. doi:10.1016/j.tree.2006.09.003.

Pardini AT, Jones CS, Noble LR, Kreiser B, Malcolm H, Bruce BD, Stevens JD, Cliff G, Scholl MC, Francis M, Duffy CAJ, Martin AP. Sex-biased dispersal of great white sharks. Nature. 2001;412(6843): 139–140.

Preston T. Who’s the real killer?. E Magazine: The Enviro Mag. 1995;6(6): 18–19.

Sminkey TR, Musick JA. Age and growth of the sandbar shark Carcharinus plumbeus, before and after population depletion. Copeia. 1995;4: 871–883.

Sminkey TR, Musick JA. Demographic analysis of the sandbar shark, Carcharhinus plumbeus, in the Western North atlantic. Fish Bull. 1996;94(2): 341–347.

Smith SE, Au DW, Show C. Intrinsic rebound potential of 26 species of Pacific sharks. Mar and Fresh Res. 1998;49(7): 663–678.

Ussami LHF. Análise da variabilidade e estruturação genética do tubarão-azul, Prionace glauca (Chondrichthyes, Carcharhinidae) na costa brasileira, utilizando marcadores microssatélites. M.Sc. Thesis, Universidade Estadual Paulista – UNESP: Instituto de Biociências de Botucatu, São Paulo. 1995. Available from: https://repositorio.unesp.br/handle/11449/99431?show=full

Weng K, Boustany A, Pyle P, Anderson S, Brown A, Block B. Migration and habitat of white sharks (Carcharodon carcharias) in the eastern Pacific Ocean. Mar Biol. 2007;152: 877–894.

